# CEACAM1 regulates the IL-6 mediated fever response to LPS through the RP105 receptor in murine monocytes

**DOI:** 10.1101/394577

**Authors:** Zhifang Zhang, Deirdre La Placa, Tung Nguyen, Maciej Kujawski, Keith Le, Lin Li, John E. Shively

## Abstract

Systemic inflammation and the fever response to pathogens are coordinately regulated by IL-6 and IL-1β. We previously showed that CEACAM1 regulates the LPS driven expression of IL-1β in murine neutrophils through its ITIM receptor. We now show that the prompt secretion of IL-6 in response to LPS is regulated by CEACAM1 expression on bone marrow monocytes. *Ceacam1^-/-^* mice over-produce IL-6 in response to an i.p. LPS challenge, resulting in prolonged surface temperature depression and overt diarrhea compared to their wild type counterparts. Intraperitoneal injection of a ^64^Cu-labeled LPS, PET imaging agent shows confined localization to the peritoneal cavity, and fluorescent labeled LPS is taken up by myeloid splenocytes and muscle endothelial cells. While bone marrow monocytes and their progenitors (CD11b^+^Ly6G^-^) express IL-6 in the early response (<2 hours) to LPS in vitro, these cells are not detected in the bone marrow after in vivo LPS treatment due to their rapid and complete mobilization to the periphery. Notably, tissue macrophages are not involved in the early IL-6 response to LPS. In contrast to human monocytes, TLR4 is not expressed on murine bone marrow monocytes. Instead, the alternative LPS receptor RP105 is expressed and recruits MD1, CD14, Src, VAV1 and β-actin in response to LPS to produce IL-6. CEACAM1 negatively regulates RP105 signaling in monocytes by recruitment of SHP-1, resulting in the sequestration of pVAV1 and β-actin from RP105. This novel pathway and regulation of IL-6 producing by CEACAM1 defines a novel role for monocytes in the fever of mice to LPS.

**AUTHOR SUMMARY:** Fever is one of the most common signs of the immune response to pathogens. The fever response to LPS or endotoxin of gram-negative bacteria is mediated by the combined action of two cytokines, IL-1β and IL-6. Regulation of their production in response to LPS is an important area of investigation. While we previously showed that the regulation of IL-1β production in neutrophils is through the lymphocyte receptor CEACAM1, we were interested if a similar mechanism operated for IL-6. Using a mouse model in which the CEACAM1 gene was knocked out, we show that IL-6 is over-produced compared to normal mice, and that monocytes, rather than neutrophils were the principal IL-6 producing cells. Surprisingly, murine monocytes do not express TLR4, the most commonly studied receptor for LPS, but instead express the low affinity LPS receptor, RP105, a receptor common expressed on B-cells. Furthermore, we show that bone marrow monocytes are rapidly released into the blood and home to tissues throughout the body in response to LPS. These findings explain much of the confusion in the literature concerning the immediate source of IL-6 and the distinct differences between murine and human monocytes in their in responses to LPS.

## INTRODUCTION

IL-6 is a central mediator of inflammation in response to a wide variety of stimuli including infection, stress and trauma ^1^. Its receptor, IL-6R, is widely expressed, especially in the liver leading to the acute phase protein response ^1^, in the hypothalamus together with IL-1β leading to systemic fever ^2^, and in the gut leading to Th17 activation ^3^. Chronic high levels of IL-6 are associated with aging, cancer, rheumatoid arthritis, neurodegenerative diseases, postmenopausal osteoporosis ^4^, and psoriasis ^5^ to name a few pathogenic conditions. As a result, activation of the IL-6 gene is under tight control, and an understanding of its regulation is fundamental to preventing a wide range of pathologies ^6^. We are especially interested in the regulation of the fever response that requires production of both IL-1β and IL-6, that together stimulate the production of PGE2 in the hypothalamus leading to a drop in the core temperature of a few tenths of a degree ^7^. Even more pronounced is the lowering of skin temperature that is perceived as “chills” followed by stimulation of the skeletal muscles or “shivering.” The critical role of IL-6 in the fever response is exemplified in IL-6 KO mice that do not exhibit a fever in response to classical stimuli such as bacterial endotoxin, viruses, and the inflammatory cytokines TNFα and IL-1β ^8^.

Given the wide range of stimuli that elicit the IL-6 response from essentially anywhere in the body, knowledge of the cells and mechanism of IL-6 secretion is essential. However, more is known about the cells that respond to IL-6 than those that produce it, and its regulation remains an area of intense investigation. We hypothesize that this regulation must be widespread, especially at the interface between the epithelium where infections, trauma and stress are likely to occur, and the immune system. A candidate gene for this regulation is CEACAM1, a homotypic cell-cell adhesion molecule ubiquitously expressed in the epithelium, constitutively expressed in neutrophils, the most abundant leukocytes, and inducibly expressed in activated lymphocytes ^9^. CEACAM1 has tissue specific, differential expression of mRNA slice forms, with an ITIM containing signaling domain expressed in the immune system and a shortened signaling domain lacking an ITIM in uninflamed epithelial cells ^10^. Notably, the expression of the ITIM containing CEACAM1 splice form is strictly regulated in response to IFNγ via IRF-1 ^11^. In agreement with this role for CEACAM1, we have previously shown that CEACAM1 regulates granulopoiesis and the systemic response to *Listeria monocytogenes* infection via the G-CSFR-STAT3 pathway ^12^, and the IL-1β response to LPS in neutrophils by a TLR4-Syk pathway ^13^. In both cases, CEACAM1 is recruited to an activated receptor (G-CSFR or TLR4), that when phosphorylated by a Src kinase on its ITIM, recruits SHP-1, which in turn, dephosphorylates the activated receptor. This is a general mechanism for CEACAM1 that has been implicated in the regulation of the insulin receptor in the liver ^14^, the EGFR in epithelial cells ^15^, and the BCR in B-cells ^16^, ^17^. In this way, CEACAM1 can moderate the effect of the immune system on stimulated epithelial cells, and when absent, as in many cancers ^18^, ^19^, the result is chronic or exaggerated inflammation. The digestive tract, including the small and large intestine, and the liver, have the highest levels of CEACAM1 expression ^20^. Since it is well known that LPS in the peritoneal cavity, mimicking leaky gut, leads to a rapid inflammatory and fever response ^21^ due to the combined actions of IL-6 and Il-1β, we speculated that an exaggerated response would be seen in *CEACAM1^-/-^* mice, providing a model system to track down the cells responsible for IL-6 release.

The plasma levels of IL-6 in *Ceacam1^-/-^* mice in response to i.p. LPS were more than twice the amount of wild type mice at 24-48 hours, including the depression of body surface temperatures and overt diarrhea in 50% of the *Ceacam1^-/-^* mice compared to none in the wild type controls. PET image analysis of mice injected i.p. with ^64^Cu-labeled-LPS exhibited LPS localization confined to the peritoneal cavity, while i.p. injection of fluorescent tagged LPS demonstrated staining in the spleen, lymph nodes and endothelial cells of skeletal muscle. Analysis of bone marrow cells revealed that a subset of bone marrow myeloid cells were rapidly mobilized to the spleen, perhaps explaining the controversy over the lack of IL-6 secreting myeloid cells in mice treated with LPS. In vitro analysis revealed that bone marrow monocytes and their progenitors produce IL-6 in the early response (<2 hours) to LPS while tissue macrophages do not. Unexpectedly, we found that TLR4, the prototypic LPS receptor of murine macrophages ^22, 23, 24^ and human monocytes and macrophages ^25, 26^ was not expressed on murine bone marrow monocytes. Instead, the alternate LPS receptor RP105, highly expressed on B-cells, was responsible for IL-6 secretion on murine bone marrow monocytes. We demonstrated that MD1, CD14, Src, VAV1 and β-actin are involved in the signaling of RP105 and that CEACAM1 regulates RP105 signaling through recruitment of SHP-1 and sequestration of pVAV1 and β-actin from pRP105 to control IL-6 producing. We conclude that CEACAM1 negatively regulates IL-6 producing in the early phase response to LPS through the RP105 signaling pathway in murine monocytes, thus defining a novel role of CEACAM1 for monocytes in the fever response.

## RESULTS

### Genetic ablation of CEACAM1 leads to an exaggerated IL-6 response to LPS

We previously showed that CEACAM1 regulates IL-1β production in LPS treated granulocytes in a TLR4-Syk specific manner ^13^. Since IL-1β and IL-6 together mediate the fever response to LPS, we performed an in vivo challenge of wild type (WT) and *Ceacam1^-/-^* mice with LPS injected i.p. Surface body temperature was measured as a sensitive indicator of the fever response along with serum multiplex cytokine levels to determine which, if any, were dysregulated in *Ceacam1^-/-^* mice treated with LPS. Phenotypically, both WT and *Ceacam1^-/-^* mice had depressed surface body temperatures, with the depression in *Ceacam1^-/-^* mice significantly lower than in WT mice at both the 8hr and 24hr time points (**Fig 1A**). 53% *Ceacam1^-/-^* mice (9 out of 17) developed overt diarrhea in comparison with none in wild type mice (**Fig 1B**). Comparison of the serum levels of cytokines between *Ceacam1^-/-^* and WT mice, revealed similar kinetics and levels for IL-1β, TNFα and IFNγ, as well as others not shown, while IL-6 levels were significantly elevated in *Ceacam1^-/-^* mice over 24 hours, returning to baseline by 48 hours (**Fig 1C-F**). The results suggest that abrogation of CEACAM1 expression in mice dramatically increases their sensitivity to i.p. LPS by specific over-expression of IL-6.

**Fig 1.**
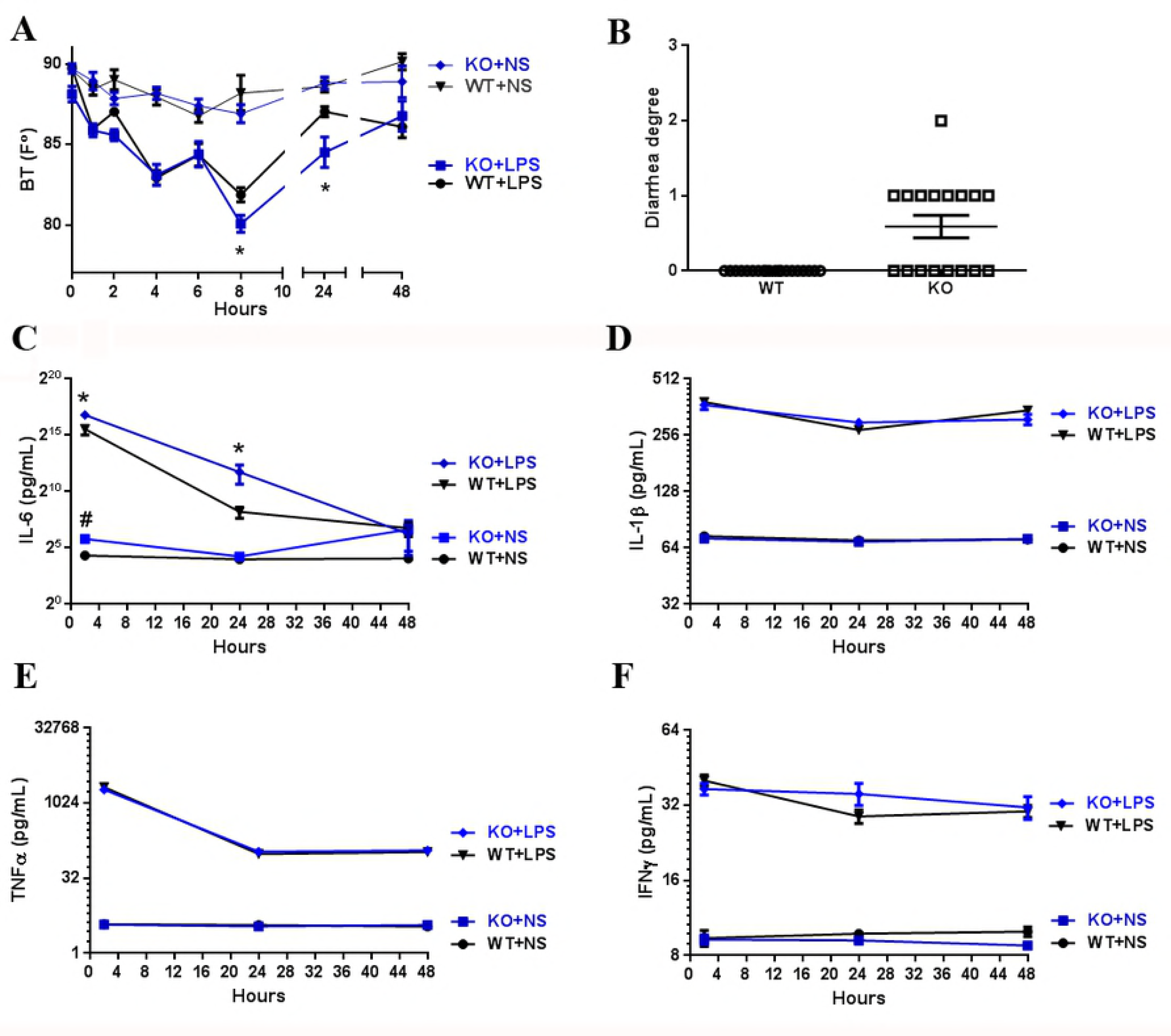
Genetic abrogation of CEACAM1 leads to decreased body surface temperature and increased diarrhea and IL-6 production in response to LPS. (A) Body surface temperature of WT and *Ceacam1^-/-^* mice in response to LPS (i.p. injection, 10 mg/kg) (n=10 each group). * p<0.05 in comparison with WT treated with LPS. # p<0.05 in comparison with WT treated with normal saline. (B) Diarrheogenic activity of *Ceacam1^-/-^* mice in response to LPS (i.p. injection, 10 mg/kg) (n=17, each group). (C-F) Quantification of 4 serum cytokines of Wild type (WT) and *Ceacam1^-/-^* mice (KO) in response to LPS (i.p. injection, 10 mg/kg) (n=10, each group). * p<0.05 in comparison with WT.

### LPS distribution after i.p. injection and IL-6 mRNA expression in mouse organs

IL-6 is considered the critical proinflammatory cytokine for the febrile response, since neither *IL-6* knock-out mice, nor animals treated with IL-6 antiserum develop fever upon peripheral immune stimulation ^8, 27, 28^. Furthermore, it is understood that IL-6 acts in concert with IL-1β as an endogenous pyrogen during LPS-induced fever ^7, 27^. Although IL-6 is reported to be synthesized and secreted by many cell types, for example, monocytes and macrophages ^29, 30^, fibroblasts ^31^, brain endothelial cells ^32, 33^, muscle cells ^34^, hepatocytes ^35, 36^, adipocytes ^37^, neurons ^38, 39^, microglial cells ^40, 41^ and astrocytes ^42, 43^, the source of serum IL-6 after i.p. treatment of LPS remains controversial ^44^. As a first approach to determining the source of systemic production of IL-6, we injected ^64^Cu-labeled LPS i.p. into mice and performed PET imaging (**Fig 2A**). This approach allows a quantitative measure of LPS localization over time. The results demonstrate that, excluding bladder secretion, ^64^Cu-DOTH-LPS is mainly localized to the peritoneal cavity, including liver, kidney and thoracic lymph nodes at 1, 2, and 4 hour time points, and is largely cleared via urinary excretion by 24 hours. Notably, very little bone activity was observed, indicating that secretion of IL-6 by bone marrow cells (if any) must be indirect. Utilizing a similar chemical procedure to produce a fluorescent version of LPS, we generated FAM-labeled LPS that is considerably brighter and more stable than commercially available FITC-LPS. Thus, as a second approach to visualizing tissue targets of LPS, FAM-LPS was injected i.p. and multiple tissues collected for analysis by immunofluorescence analysis at 1 hour. The results demonstrated high uptake into the spleen, lymph nodes, and the endothelial cells of skeletal muscle (**Fig 2B, 2D**). Further analysis of the spleen cells labeled indicated that they were macrophages (**Fig 2C**).

**Fig 2.**
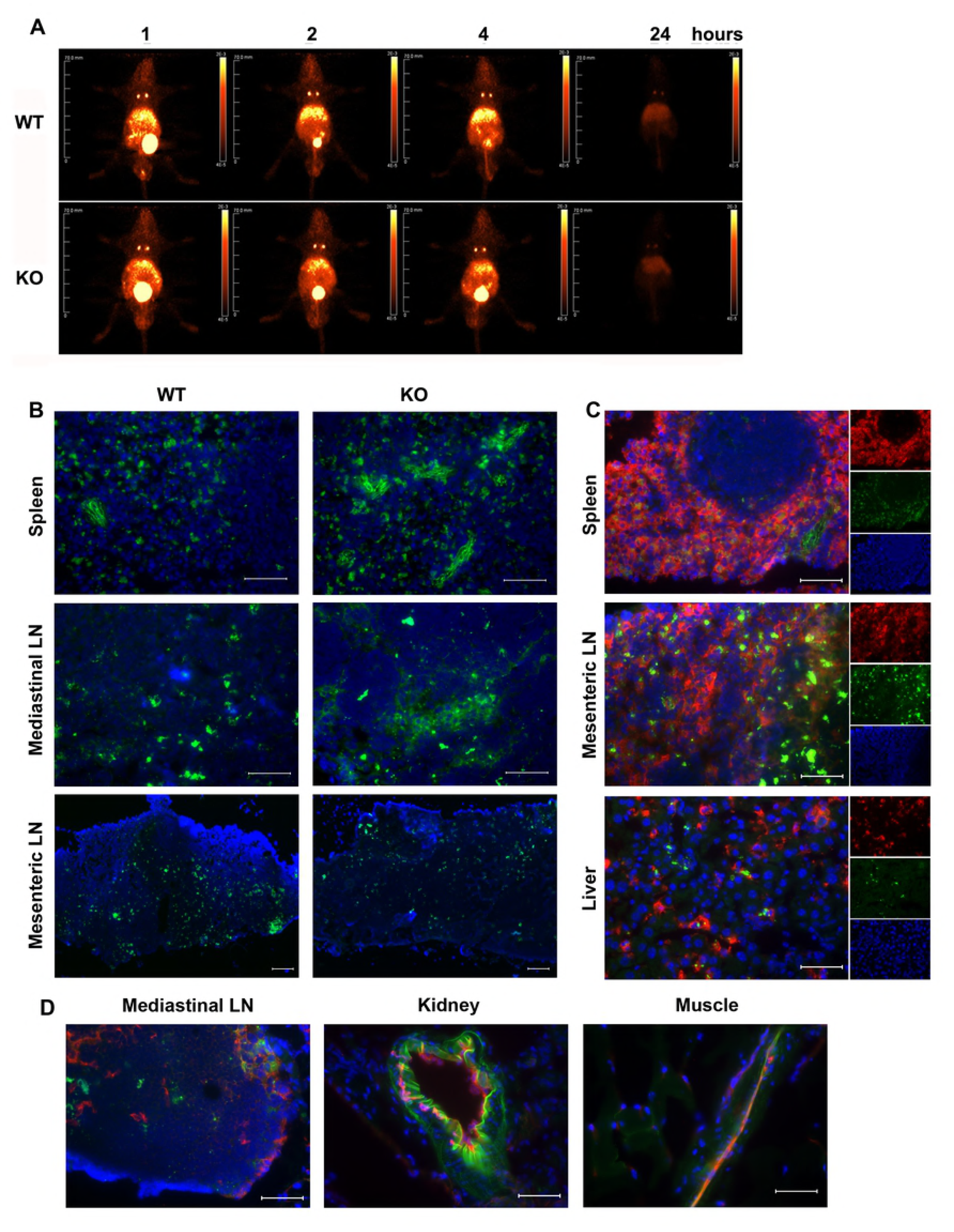
Distribution of intraperitoneal injection of ^64^Cu-labeled LPS or FAM-LPS. (A) PET imaging of i.p. injection of ^64^Cu-labeled LPS over time. (B-D) Immunofluorescent staining of selected tissues 1 hour after i.p. injection of FAM-LPS.

Since the first two approaches only indicate tissues of LPS uptake and not IL-6 production, we also measured *IL-6* mRNA by qPCR of peritoneal tissues and other organs. Most peritoneal cavity tissues/organs including mesentery, peritoneal membranes, pancreas, and fatty tissues did not show any difference between WT and *Ceacam1^-/-^* mice, while omentum and small intestine exhibited a decrease in *Ceacam1^-/-^* mice (**Fig S1**). Surprisingly, skeletal muscle, lung, and kidney also exhibited significant decreases in the *IL-6* mRNA expression compared to WT counterparts, while brain, bone marrow cells and mesenteric lymph nodes had no difference. The organs with significantly increased levels of *IL-6* mRNA in Ceacam1^-/-^mice in comparison with WT mice were liver and spleen (**Fig 3A**). Given the large size of the liver, the tentative conclusion is that liver may be the main IL-6 producer in response to i.p. LPS, followed by the spleen.

**Fig 3.**
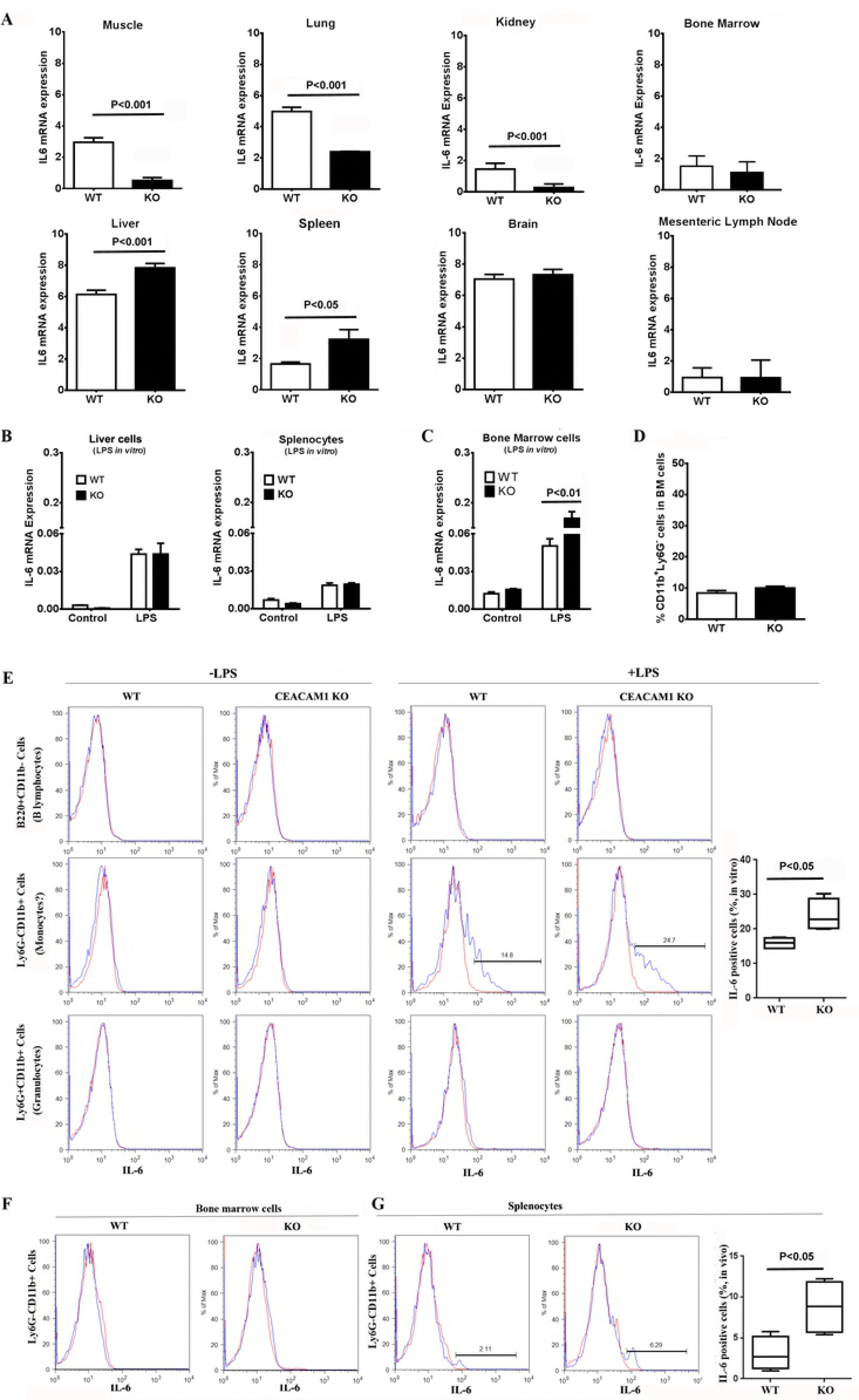
IL-6 expression levels of organs in *Ceacam1^-/-^* mice in response to LPS challenge. (A) *IL-6* mRNA expression levels of mouse organs after i.p. injection of LPS (10mg/kg) for 2 hours (n=4). (B) *IL-6* mRNA expression level in PBS perfused liver and splenocytes after treated with 500 ng/mL LPS for 2 hours in vitro (n=3). (C) IL-6 mRNA expression level in bone marrow cells after treated with 500 ng/mL LPS for 2 hours in vitro (n=3). (D) Percentage of bone marrow CD11b^+^Ly6G^-^ cells in WT and *Ceacam1^-/-^* mice (n=4). (E) Intracellular staining of IL-6 in bone marrow cells after treated with Brefeldin A (BFA) plus 500ng/mL LPS for 5 hours in vitro (n=3). (F) Intracellular staining of IL-6 of bone marrow cells after i.p. injection of BFA plus LPS (10mg/kg) for 5 hours in vivo (n=3). (G) Intracellular staining of IL-6 of splenocytes after i.p. injection of BFA plus LPS (10mg/kg) for 5 hours in vivo (n=3).

### Liver cells and splenocytes are not the source of IL-6 in early response to LPS

To explore the role of liver and spleen in the secretion of IL-6 in response to LPS, liver cells (from PBS perfused liver to remove blood cells) and splenocytes from untreated mice to determine which cells, if any, produce IL-6 in direct response to LPS in vitro. Our results showed that IL-6 mRNA expression of liver cells and splenocytes after treated with LPS for 2 hours were not different between WT and *Ceacam1^-/-^* mice (**Fig 3B).** When isolated hepatocytes or Kupffer cells were treated with LPS plus BFA (brefeldin A) for 5 hours, hepatocytes from *Ceacam1^-/-^* mice were negative for both IL-6 and TNFα (**Fig S2A**) while Kupffer cells were positive for TNFα only **(Fig S2B)**. Furthermore, the IL-6 secretion of liver cells in response to LPS treatment was not significant difference between the two mouse strains until after 6 hours (**Fig S3A**).

Based on these results, we surmised that IL-6 production observed for the liver (**Fig 3A**) was due to blood cells trapped at the time of sacrifice. Therefore, the analysis was repeated on liver perfused after i.p. LPS treatment. There were no differences in *IL-6* mRNA levels between WT and *Ceacam1^-/-^* mice (data not shown), suggesting that blood cells trapped in the liver, rather than endogenous cells were responsible for the observed difference (**Fig 3A**). Western blot analysis of phospho-gp130, the key activation signal transducer of the IL-6 receptor, revealed that livers of both WT and *Ceacam1^-/-^* mice had similar levels after LPS treatment (**Fig S3B**). The same was true for the downstream effectors of the gp-130, pSTAT1, pSTAT3 and SOCS3 (**Fig S3B**). We conclude that although the liver is a major organ responsive to IL-6, it is not the main source of IL-6 in the early response to LPS.

### A subgroup of bone marrow CD11b^+^Ly6G^-^ myeloid cells secrete IL-6 and are mobilized in the early response to LPS

Since it was likely that the source of the IL-6 producing cells in the liver and spleen originated from the bone marrow, we collected bone marrow cells from untreated mice and determined their in vitro production of IL-6 in response to LPS. This analysis revealed significantly higher levels of *IL-6* mRNA for *Ceacam1^-/-^* vs WT bone marrow cells in response to LPS (**Fig 3C**).

Analysis of cell surface markers of bone marrow cells together with intracellular IL-6 staining after LPS treatment revealed that a subgroup of CD11b^+^Ly6G^-^cells but not CD11b^+^Ly6G^+^ cells (granulocytes) or B220^+^ cells (B lymphocytes) produced IL-6 (**Fig 3E**). Notably, there was no significant difference in CD11b^+^Ly6G^-^ cell percentages of bone marrow cells between WT and *Ceacam1^-/-^* mice (**Fig 3D**), suggesting that the number of IL-6 producing cells in the bone marrow per se are not responsible for the IL-6 differences observed in between WT and *Ceacam1^-/-^* mice. In accordance with the negative finding of *IL-6* mRNA in bone marrow cells treated with LPS in vivo (**Fig 3A**), intracellular staining of IL-6 was negative in bone marrow cells after i.p. injection of LPS plus BFA (**Fig 3F**), but positive in one subgroup of CD11b^+^Ly6G^-^ cells in the spleen (**Fig 3G**). These results suggest that IL-6 producing bone marrow cells were mobilized from the bone marrow to the spleen after i.p. LPS treatment and that the subgroup of CD11b^+^Ly6G^-^ cells may be responsible for the difference of IL-6 production between WT and *Ceacam1^-/-^* mice after LPS challenge.

### Monocytes and the progenitors of myeloid CD11b^+^Ly6G^-^ cells are IL-6 producing cells in the early response to LPS

Since CD11b^+^Ly6G^-^ cells in the bone marrow include different groups of myeloid cells, further analysis of cell surface markers was performed to clarify which cell type was responsible for IL-6 production. Recent studies have characterized bone marrow CD115 (M-CSF-R) positive cells into monocytes (Mo), common monocyte progenitors (cMoP), monocyte-macrophage DC progenitors (MDP) and common DC precursor (CDP) according to cell surface markers CD117 (c-Kit) and CD135 (FLT-3) ^45^. Cell surface staining showed that all of four populations are CEACAM1 positive in WT mice **(Fig S4A)**.

After in vitro treatment of bone marrow cells with LPS, CD115 expression was down-regulated while the staining pattern of MDP, cMoP, Mo and CDP were indistinguishable (**Fig S4B**). Therefore, the four populations of MDPs (Lin^-^CD115^+^CD117^+^CD135^+^), cMoP (Lin^-^ CD115^+^CD117^+^CD135^-^), Mo (Lin^-^CD115^+^CD117^-^CD135^-^), and CDP (Lin^-^CD115^+^CD117^-^ CD135^+^) were sorted (**Fig 4A**) and treated with LPS plus BFA for 5 hours. In WT mice, cMoP and Mo but not MDP and CDP were IL-6 positive. Surprisingly, in *Ceacam1^-/-^* mice, cMoP, Mo, and MDP but not CDP were all positive for IL-6 with significantly increased percentages over WT (**Fig 4E-F**). There were no significant differences in the percentages of Lin^-^ cells (**Fig 4B**), CD115^+^ cells (**Fig 4C**), nor the MDP, cMoP, Mo, and CDP subsets between WT and *Ceacam1^-/-^* mice (**Fig 4D**). These analyses show that monocytes and their progenitors are the major IL-6 producing cells in the bone marrow in the early response to LPS, and that the absence of CEACAM1 results in high levels of IL-6 production.

**Fig 4.**
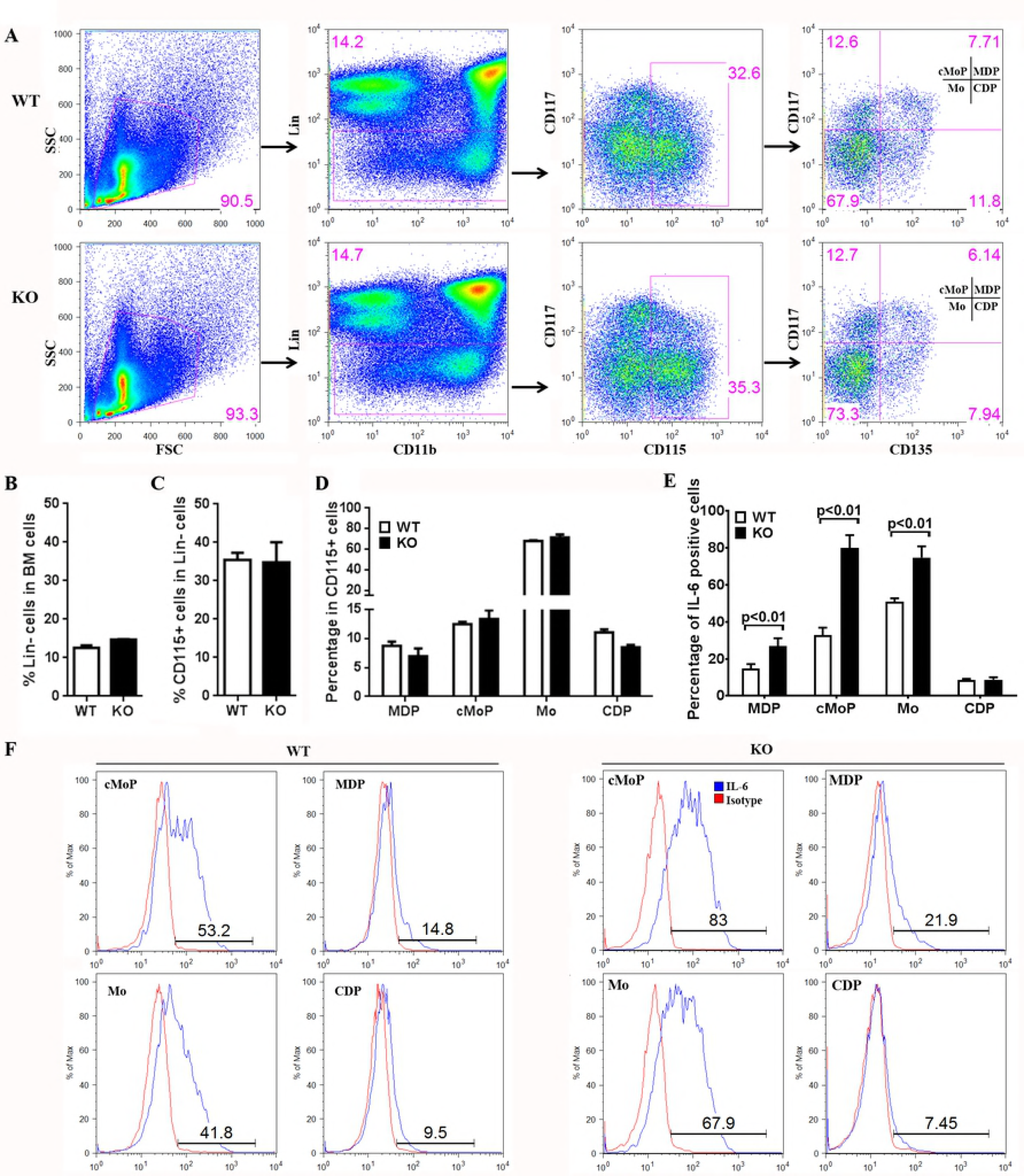
Production of IL-6 in monocytes and their progenitors in wild type and *Ceacam1 ^-/-^* mice. (A) Flow cytometry gating of monocytes (Mo), common monocyte progenitors (cMoP), monocyte-macrophage DC progenitors (MDP) and common DC precursors (CDP) in bone marrow cells (n=4). (B) Percentage of Lin^-^ cells in bone marrow cells (n=4). (Note: Lin = lineage markers: including CD3, CD19, B220, Ly6G, NK1.1, TER119) (C) Percentage of CD115^+^ cells in Lin^-^ bone marrow cells (n=4). (D) Percentage of MDP, cMoP, Mo, and CDP in Lin^-^CD115^+^ bone marrow cells (n=4). (E) Percentage of intracellular IL-6 staining in sorted MDP, cMoP, Mo, and CDP cells after treatment with Brefeldin A (BFA) plus 500ng/mL LPS for 5 hours in vitro (n=4). (F) Histogram of intracellular IL-6 staining of sorted MDP, cMoP, Mo, and CDP cells after treatment with BFA plus 500ng/mL LPS for 5 hours in vitro (n=4).

### Macrophages do not produce IL-6 in the early response to LPS

Since monocytes and macrophages differ in their ability to process pro-IL-1β and release mature IL-1β ^46, 47^, it was necessary to determine if a similar situation occurred for IL-6 production and secretion in macrophages. Peritoneal cavity macrophages, isolated from WT and *Ceacam1^-/-^* mice, were treated in vitro with LPS for 2, 4 and 24 hours. The results showed that there is no difference in *IL-6* mRNA levels between WT and *Ceacam1^-/-^* mice at the 2 hour time point, but *IL-6* mRNA levels significantly increased at the 4 hour time point and decreased at the 24 hour time point in *Ceacam1^-/-^* mice (**Fig S5A**). When splenocytes from *Ceacam1^-/-^* mice were treated with LPS plus BFA for 5 hours, all three groups of myeloid cells (Ly6G^-^ CD11b^+^ cells), granulocytes (Ly6G^+^CD11b^+^ cells), and lymphocytes (Ly6G^-^CD11b^-^ cells) were negative for IL-6 by intracellular staining (**Fig S5B**). Similar results were obtained for WT mice (data not shown). Furthermore, the murine macrophage cell line RAW264.7 treated with LPS plus BFA for 5 hours was negative for intracellular IL-6 production (**Fig S5C**). In fact, RAW264.7 cells treated with LPS showed that *TNFα* rather than *IL-6* mRNA significantly increased at 2 hours in comparison with untreated controls, and that *IL-6* mRNA levels only started to increase at the 4 hour time point, maintaining the increase through the 24 hour time point (**Fig S5D**). The release of IL-6 into the supernatant after LPS treatment was delayed until the 24 hour time point but not at the early 2 hour time point (**Fig S5E**). Intracellular IL-6 staining analysis showed that these cells began to produce IL-6 only after 11 hours of LPS treatment (**Fig S6A**). Furthermore, silencing of CEACAM1 with siRNA in RAW264.7 cells did not affect IL-6 secretion at the 2 hour time point (**Fig S6B**). Another murine macrophage cell line, J774A.1, exhibited similar results as RAW264.7 cells, i.e., there was no IL-6 production within 5 hours after LPS treatment and CEACAM1 siRNA did not interfere with IL-6 production (**Fig S7**). Moreover, Kupffer cells in the liver (analyzed above, **Fig S2B**) gave similar results as did spleen macrophages (**Fig S5B)**.These data indicate that although macrophages are derived from either monocytes or the yolk sac and are self-replenished ^48^, they are unable to synthesize and release IL-6 in the early phase (<2 hours) of the LPS response.

To further explore the early response to LPS treatment, the time course of *IL-6* mRNA was measured in bone marrow monocytes. As shown in **Fig 5A**, *IL-6* mRNA is significantly increased as early as 30 minutes after LPS treatment and reached a peak at 90 min in *Ceacam1^-/-^* mice. These data confirm that monocytes are the main source of IL-6 in the early response to LPS and that CEACAM1 regulates this response. Given the report that an IL-6 feed-forward loop can increase IL-6 production ^49, 50^, *Il6ra* ^-/-^ mice and *Stat3^flox/flox^* mice were included in our study. The analysis of *IL-6* mRNA expression of bone marrow monocytes from *Il6ra* ^-/-^ mice (**Fig 5A**) and *Stat3^flox/flox^* mice (**Fig 5B**) demonstrated that a different expression pattern compared to the *Ceacam1*^-/-^ mice over the time course of LPS treatment. These results suggest CEACAM1 expression does not interfere with the IL-6 receptor signal pathway, but only in the response to LPS.

**Fig 5.**
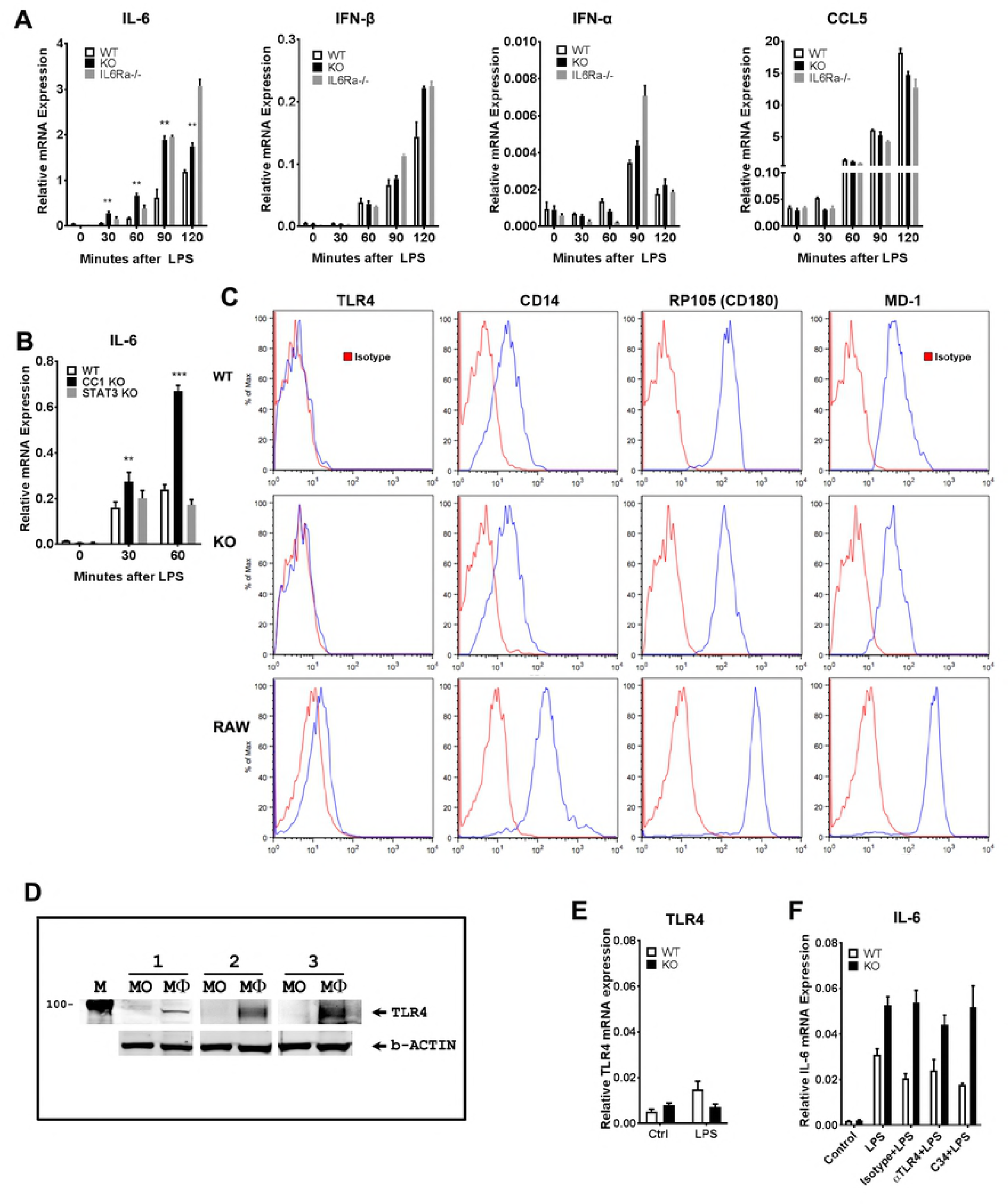
Cytokine expression of bone marrow monocytes. RP105 and not TLR4 is expressed. (A) *IL-6, IFN-β, IFN-α* and *CCL5* mRNA expression levels of bone marrow monocytes in WT, *Ceacam1^-/-^* and *Il6ra^-/-^* mice after treatment of 500ng/mL LPS in vitro (n=3). (B) *IL-6* mRNA expression levels of bone marrow monocytes in WT, *Ceacam1^-/-^* and *Stat3^flox/flox^* (*Stat3^-/-^*) mice after treatment of 500ng/mL LPS in vitro (n=3). (C) Surface TLR4, CD14, RP105 and MD1 staining of RAW264.7 cells and bone marrow monocytes of WT and *Ceacam1^-/-^* mice (n=3). (D) Immunoblot analysis for TLR4 detection in RAW264.7 cells and bone marrow monocytes of WT mice (1, 2, 3 mean three separated experiments; MO: monocytes; Mϕ: RAW264.7 macrophages). (E) *TLR4* mRNA expression level in bone marrow monocytes of WT and *Ceacam1^-/-^* mice(n=4). (F) *IL-6* mRNA expression level in bone marrow monocytes pretreated with TLR4 blocking antibody (10μg/mL) or TLR4 inhibitor C34 (100μM) for 20 minutes following 500ng/mL LPS treatment for 30 minutes in vitro (n=3).

### TLR4 is not expressed on murine bone marrow monocytes

Having identified bone marrow monocytes as the source of prompt IL-6 secretion in response to LPS, we proceded to analyze the mechanism of CEACAM1 regulation. Since our previous studies showed CEACAM1 regulated TLR4 signaling in murine neutrophils, we expected a similar mechanism in murine monocytes, especially since human monocytes express abundant amounts of TLR4 ^26^. TLR4, the canonical LPS receptor, signals through the MyD88- and Toll/IL-1R domain-containing adapter inducing IFN-β (TRIF)-dependent upstream signals that lead to the production of proinflammatory (IL-6 and IL-1β) and anti-inflammatory mediators (IFN-β, IFN-α, CCL5), respectively ^51, 52^. However, unlike *IL-6* mRNA production in LPS treated murine monocytes, there was no change in the levels of *IFN-β, IFN-α,* and *CCL5* mRNA at the 30 minute time point in WT, *Ceacam1*^-/-^, and *Il6ra* ^-/-^ mice. *IFN-β, IFN-α,*and *CCL5* mRNA levels began to increase after 60 minutes but did not exhibit differences between WT and *Ceacam1*^-/-^ mice at both the 60 and 90 minute time points. These results suggests that the TRIF-dependent signaling pathway, as one of two main signaling pathways of TLR4, was not directly involved in the response of murine bone marrow monocytes to LPS. Furthermore, TLR4 expression was negative on bone marrow monocytes from both WT and *Ceacam1^-/-^* mice using anti-TLR4 antibody surface staining (**Fig 5C**) and western blot analysis (**Fig 5D**) even though *TLR4* mRNA was detected at low levels by qPCR (**Fig 5E**). When RAW264.7 macrophage lineage cells were used as a positive control, TLR4 was easily detected by both surface staining and western blot analysis. On the other hand, CD14 that acts as an LPS co-receptor for TLR4, was strongly positive for bone marrow monocytes (**Fig 5C**). We also performed intracellular immunofluorescent staining to explore the possibility that TLR4 was localized to intracellular granules, but the monocytes were negative (data not shown). In addition, TLR4 blocking antibody and the TLR4 inhibitor C34 did not interfere with LPS-induced *IL-6* mRNA expression in these cells treated with LPS (**Fig 5F**). Taken together, we conclude that TLR4 is not expressed on murine bone marrow monocytes and that TLR4 signaling is not responsible for the observed IL-6 production of murine monocytes.

### RP105 (CD180) on bone marrow monocytes is the LPS receptor responsible for the early IL-6 response to LPS

In consideration of alternative receptors for LPS, it is well known that B-cells respond strongly to LPS ^53^. B-cells have two distinct LPS receptor complexes, TLR4/MD2 and RP105/MD1 ^54^. The extracellular domains of TLR4 and RP105 associate with MD2 and MD1, respectively, to form heterodimers, thereby forming the binding sites to LPS ^54, 55^. Although we did not detect TLR4 on murine monocytes, RP105 and MD1 were highly expressed on bone marrow monocytes from both WT and *Ceacam1^-/-^* mice (**Fig 5C**). Since the signaling pathway for RP105 has been extensively studied and involves recruitment of VAV1 and β-actin ^54, 56^, we performed a number of co-IP studies on murine bone marrow monocytes treated with LPS. MD1 co-IPed with RP105 in the presence or absence of LPS in bone marrow monocytes (**Fig 6A**), while RP105 co-IPed with pVAV1 and β-actin in the absence of LPS in WT and *Ceacam1^-/-^* mice (**Fig 6B).** After treatment with LPS, pVAV1 and β-actin dissociated with RP105 in WT mice, but in *Ceacam1^-/-^* mice, pVAV1 and β-actin remained associated with RP105. IP of CEACAM1 in WT mice co-IPed β-actin after LPS treatment (**Fig 6C**). These results suggest that CEACAM1 can sequester pVAV1 and β-actin from RP105 after LPS treatment thus negatively regulating RP105 downstream signaling. In the absence of CEACAM1, RP105 remains associated with pVAV1 and β-actin resulting in increased downstream signaling. Furthermore, the VAV1 inhibitor azathioprine, and a metabolite of azathioprine, 6-thio-GTP, was able to block the *IL-6* mRNA over-response to LPS in *Ceacam1^-/-^* mice (**Fig 6D**). Moreover, an RP105 activating monoclonal antibody ^57^ was able to stimulate *IL-6* mRNA expression in WT mice and *IL-6* mRNA over-expression in *Ceacam1^-/-^* mice similar to LPS treatment (**Fig 6E**). On the other hand, blocking antibodies to MD1 or CD14 completely abrogated LPS-induced IL-6 mRNA expression in both WT and *Ceacam1^-/-^* mice (**Fig 6F**). Furthermore, CD14 was co-IPed with RP105 (**Fig 6G**). We conclude that the RP105/MD1/CD14 complex on murine bone marrow monocytes is responsible for the early phase expression of IL-6 in LPS treated mice, and CEACAM1 negatively regulates RP105 signaling in response to LPS stimulation.

**Fig 6.**
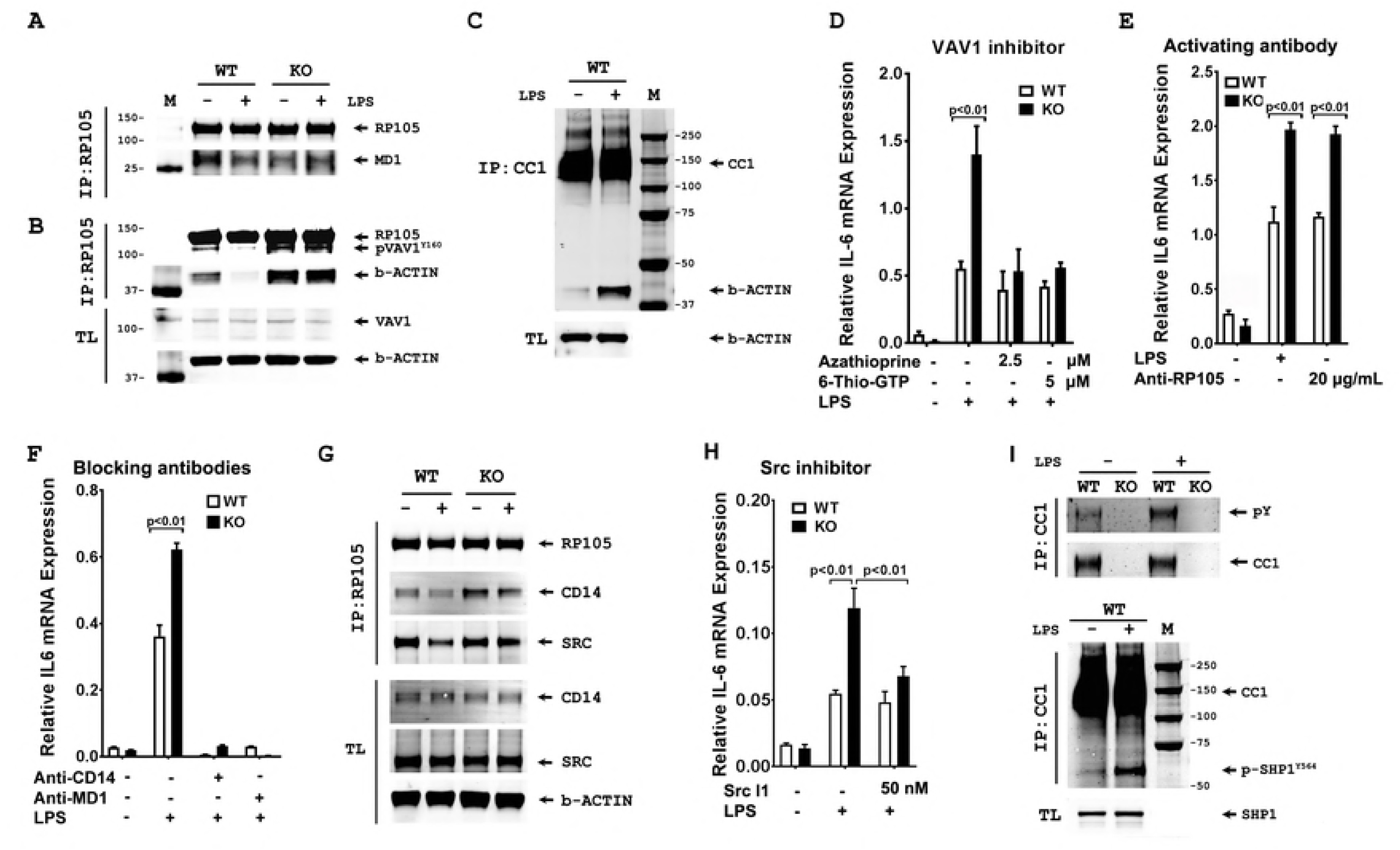
Co-immunoprecipitation and immunoblot analyses of RP105 and its signaling partners on bone marrow monocytes in response to LPS in Ceacam1^-/-^ and wild type mice. (A) Immunoblot analysis of RP105, MD1 and Syk after immunoprecipitation (IP) with RP105 antibody in WT and *Ceacam1^-/-^* bone marrow monocytes with or without 500ng/mL LPS treatment for 15 minutes. (B) Immunoblot analysis of RP105, pVAV1 and β-actin after IP with RP105 antibody in WT and *Ceacam1^-/-^* bone marrow monocytes with or without 500ng/mL LPS treatment for 15 minutes (TL: total cell lysate). (C) Immunoblot analysis of CEACAM1 and β-actin after IP with CEACAM1 antibody in WT bone marrow monocytes with or without 500ng/mL LPS treatment for 15 minutes (TL: total cell lysate). (D) *IL-6* mRNA expression of bone marrow monocytes pretreated with pVAV1 inhibitor azathioprine and 6-thio-GTP for 20 minutes following 500ng/mL LPS treatment for 30 minutes in vitro (n=3). (E) *IL-6* mRNA expression of bone marrow monocytes treated with RP105 activating antibody (clone RP/14, 20μg/mL) or 500ng/mL LPS for 30 minutes (n=3). (F) *IL-6* mRNA expression of bone marrow monocytes treated with blocking CD14 antibody (clone M14-23, 10μg/mL) or blocking MD1 antibody (clone MD113, 10μg/mL) for 20 minutes following 500ng/mL LPS for 30 minutes (n=3). (G) Immunoblot analysis of CD14 and Src after IP with RP105 antibody in WT and *Ceacam1^-/-^* bone marrow monocytes with or without 500ng/mL LPS treatment for 15 minutes (TL: total cell lysate). (H) *IL-6* mRNA expression of bone marrow monocytes pretreated with 50nM Src l1 (Src inhibitor) for 20minutes following treatment of 500ng/mL LPS for 30 minutes (n=3). (I) Anti-CEACAM1 IPs from bone marrow monocytes after treatment with 500ng/mL LPS for 15 minutes were followed by immunoblot analysis with 4G10 or anti-pSHP1 antibodies. (Upper); IP with anti-CEACAM1, immunoblot analysis with 4G10. Loading control: immunoblot with anti-CEACAM1. (Lower): IP with anti-CEACAM1 and immunoblot with anti-pSHP-1. Loading control: immunoblot with anti-SHP1. (TL: total cell lysate).

It was reported that Lyn phosphorylation and its kinase activity were involved in RP105 signaling in B cells ^54^. In our study of mouse monocytes, when RP105 was IPed, we did not detect Lyn, but instead, we found RP105 co-IPed with Src (**Fig 6G).** A selective and competitive dual site Src inhibitor (Src l1) was able to block the over-response of *IL-6* mRNA expression to LPS in *Ceacam1^-/-^* mice (**Fig 6H**). This suggests that Src, but not Lyn, is the kinase involved in RP105 mediated LPS signaling in murine monocytes.

We have previously shown that CEACAM1 act as an inhibitory B-cell co-receptor through recruitment of the inhibitory tyrosine phosphatase SHP-1 ^16^. Suspecting a similar inhibitory mechanism in LPS treated monocytes, we IPed CEACAM1 in LPS treated monocytes and performed western blot analysis with anti-phosphotyrosine and anti-phospho-SHP-1 antibodies (**Fig 6I**). The results demonstrate that CEACAM1 is phosphorylated on tyrosine after LPS treatment (**Fig 6I upper**), that phospho-SHP-1 is co-IPed with CEACAM1, and the levels of phhospho-SHP1increase after treatment with LPS (**Fig 6I lower**). Furthermore, treatment of bone marrow monocytes with LPS in the presence of SHP1 inhibitor, PTP inhibitor III, increases phospho-VAV1 in WT but not in *Ceacam1^-/-^* mice mice (**Fig S8**). These results show that RP105 stimulation by LPS leads to recruitment of SHP-1 to CEACAM1, a result that limit tyrosine phosphorylation of both VAV1 and RP105 by Src. Thus, murine monocytes may share a common signaling pathway with B-cells since they both express CEACAM1 and RP105.

## DISCUSSION

IL-6 is a soluble mediator with a pleiotropic effect on inflammation, immune response and hematopoiesis affecting vascular disease, lipid metabolism, insulin resistance, mitochondrial activities, the neuroendocrine system and neuropsychological behavior ^58^. In this study, we focused on the role of CEACAM1 in the regulation of IL-6 production. Rather surprisingly, we found that bone marrow monocytes and not peripheral macrophages were responsible for the early IL-6 response to i.p. LPS. Although there was an increase of *IL-6* mRNA expression in macrophages after 2 hours, secreted IL-6 occurred much later and was likely due to the early production of TNFα by these cells. Moreover, the bone marrow monocytes responsible for IL-6 production as shown by in vitro analysis, were not detected in the bone marrow of mice treated in vivo with LPS, demonstrating their rapid mobilization to the periphery. For example, analysis of the liver of LPS treated mice revealed that the source of IL6 was in the blood rather than in isolated hepatocytes or Kupffer cells. In accordance with this finding, it was earlier reported that low concentrations of Toll-like receptor (TLR) ligands in the bloodstream drive CCR2-dependent emigration of monocytes from the bone marrow ^59^. In other results, we found that the CCR2 ligand, CCL-2 was elevated in both WT mice and *Ceacam1^-/-^* mice treated with LPS, with the higher levels in *Ceacam1^-/-^* mice (data not shown).

The discovery of TLR family proteins was particularly critical in showing the importance of innate immunity in the host defense against microbial infection. TLRs are characterized by extracellular leucine-rich repeat (LRR) motifs and intracellular Toll/interleukin 1 receptor (TIR) domains ^52^. TLR4 is a well-known pathogen recognition receptor that plays a key role in the prototypical inflammatory stimulus to LPS ^60^. LPS engagement of TLR4 initiates a cascade of signaling events via intracellular Toll/IL-1R signaling domains, that involve the primary recruitment of the Mal adaptor protein and its subsequent association with My88 to ultimately promote the activation of the NF-κB transcriptional complex and induction of many pro-inflammatory cytokine genes, such as *IL-6, TNFα, and IL-1β*. Host immune responses triggered by TLR4 also involve the recruitment of other intracellular signaling adaptors, in particular, TRIF and TRIF-related adaptor molecule, which also facilitate activation of NF-κB and IRF3 (IFN regulatory factor 3), the latter of which promotes the transcription of proinflammatory type I IFN genes ^61^. Unexpectedly, our results demonstrated that TLR4 protein is not expressed on murine bone marrow monocytes as evidenced by lack of activation of the TRIF-dependent pathway, absence of TLR4 protein and lack of effect on IL6 production by a TLR4 blocking antibody or TLR4 inhibitor C34. Although it is well known that TLR expression is high on human monocytes, many murine cells of myeloid origin, including macrophages, microglia, myeloid DCs, and granulocytes have been reported to have high levels of TLR4 expression ^62^. However, the status of TLR4 expression in murine monocytes, neither TLR4 mRNA nor TLR4 protein was shown ^63^. In contrast, mouse macrophages, as well as the two macrophage cell lines RAW264.7 and J774A.1, express TLR4 on their cell surface. However, neither of these cell lines, nor peripheral murine macrophages, exhibited an early IL-6 response to LPS (< 2 hours). Thus, most studies on LPS stimulated TLR signaling in the mouse are limited to macrophages, while in human, both monocytes and macrophages are used. A further source of confusion, relevant to our study is that mouse monocytes are CEACAM1^+^ and TLR4^-^ (our data) while human monocytes are CEACAM1^-^ and TLR4^+^(**Fig S9**). Notably, when we generated a human *Ceacam1* transgenic (TG) mouse using the complete human CEACAM1 genome ^64^ and crossed them into the *Ceacam1^-/-^* background, bone marrow monocytes in the *hCeacam1* TG mice did not express hCEACAM1 protein (data not shown).

The finding that TLR4 negative murine bone marrow monocytes were responsible for the prompt IL6 response to LPS necessitated a search for an alternative LPS receptor. The obvious candidate, RP105, was first reported in 1995 as a LRR protein expressed on B cells ^65^. Although RP105 has only 11 amino acids in the intracellular portion and lacks a TIR domain, ligation of RP105 with anti-RP105 monoclonal antibody (mAb) transmits powerful activation signals in B cells, including proliferation ^66^. RP105 shares some features with TLR4. First, RP105 is associated with MD-1, an MD-2 homolog. Second, both RP105 and TLR4 contain 22 LRRs in their extracellular portions, suggesting the possible involvement of RP105/MD-1 in the LPS-induced response. In fact, RP105-deficient mice as well as MD-1-deficient mice show reduced LPS-dependent proliferation and CD86 up-regulation in B cells, albeit to a lesser extent than TLR4-deficient mice. Third, although LPS appears to bind to MD-1 with lower affinity than to MD-2 ^67^, the RP105/MD-1 complex is expressed not only on B cells but also on macrophages and dendritic cells ^68^. We now report that the RP105/MD1 complex is also expressed on murine bone marrow monocytes and is negatively regulated by CEACAM1 as evidenced by sequestrating pVAV1 and β-actin from pRP105 in WT mice and the increased association of RP105 with pVAV1 and β-actin in CEACAM1^-/-^ mice. The involvement of pVAV1 and β-actin in LPS stimulated RP105 signaling in B-cells has been previously reported ^55^. Furthermore, we show that CEACAM1 in murine monocytes is phosphorylated on tyrosine and recruits the inhibitory tyrosine phosphatase SHP1 after treatment of with LPS, a finding similar to our previous studies on murine neutrophils treated with LPS ^12^. More importantly, CEACAM1, itself an actin recruiting receptor ^69^, competes with VAV1 for actin recruitment, thus diminishing the ability of VAV1 to signal downstream to mediators such as NFkB, required for IL-6 expression. The dramatic sequestration of actin away from VAV1 is shown in **Fig 6C**. The overall model for LPS/RP105 signaling in murine monocytes in shown in **Fig 7**.

**Fig 7.**
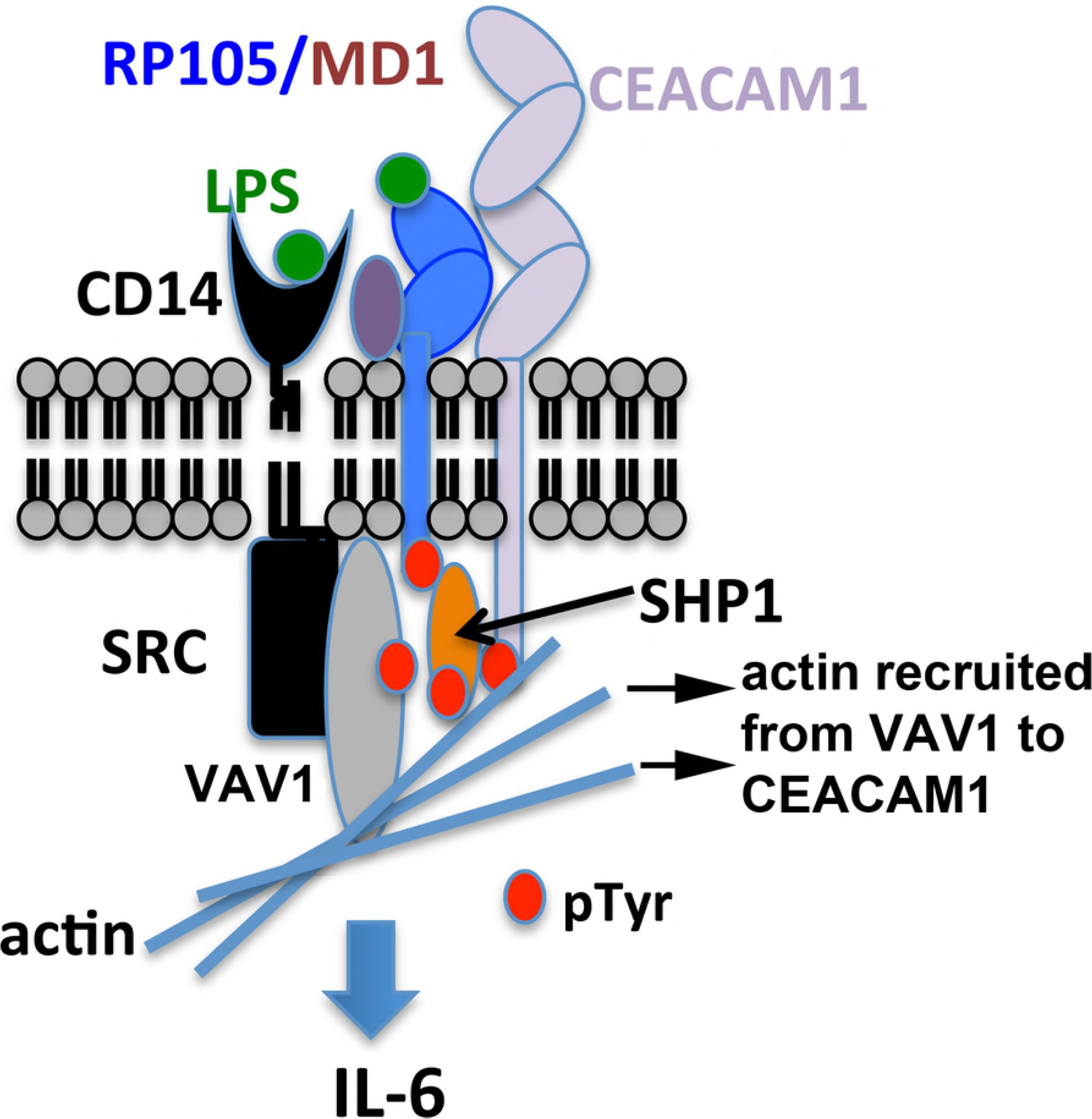
Model for regulation of RP105 signaling in LPS treated murine monocytes. RP105 (blue) associates with MD1 (purple) and CD14 (black). In response to LPS, the complex recruits SRC (black), VAV1 (grey), actin (lt. blue) and CEACAM1 (mauve). SRC phosphorylates tyrosines (red) on RP105, VAV1, and CEACAM1. SHP1 (orange) is recruited to the pITIM on CEACAM1, and in turn, is phosphorylated on its tyrosine by SRC. Downstream signaling of VAV1, dependent on recruitment of actin, is reduced by sequestration of actin by CECAM1 (shown by arrows).

It is well-known that macrophages and dendritic cells are monocyte-differentiated cells that express both TLR4/MD2 and RP105/MD1. In agreement with our finding that macrophages are more tuned to LPS/TLR4 signaling, RP105- or MD-1-deficient macrophages are not impaired in TNFα production induced by the lipid A moiety of LPS ^68^. While DCs from RP105-deficient mice produced significantly higher concentrations of proinflammatory cytokines after stimulation with LPS, the RP105/MD-1 complex competes with the binding of LPS to the TLR4-MD-1 complex and negatively regulate LPS-TLR4-mediated responses in dendritic cells ^70^. If TLR4 is a high affinity receptor for LPS while RP105 a low affinity receptor, the differential expression between macrophages/dendritic cells and murine bone marrow monocytes may be a fine-tuning mechanism to prevent an over-response to LPS, in this case, initiation of the fever response. Moreover, peripheral blood monocytes, acting as adult stem cells, are capable of undergoing maturation into several types of tissue-resident macrophages, including tissue resident macrophages, Kupffer cells, Langerhans cells of the skin, dendritic cells, microglia, and osteoclasts ^71^. In the process of their tissue differentiation their requirement for sensitivity to LPS may change.

A major finding of our study is that CEACAM1, previously shown to regulate LPS signaling in neutrophils, also regulates LPS signaling in monocytes, but in the case of neutrophils the inhibitory regulation is through TLR4 ^12^, while in monocytes through RP105. In both cases, recruitment of the inhibitory tyrosine phosphatase SHP1 is involved, suggesting that the ITIM sequence in CEACAM1 ^12^ plays a major role in dampening immune responses in a wide variety of immune cells. Indeed, that is also the case for B-cells ^16^ and T-cells ^19^. A second major finding of our study is that the prompt IL6 mediated fever response to LPS occurs through monocytes in the mouse and that it is negatively regulated by CEACAM1. Indeed, CEACAM1^-/-^ mice experience hyper IL6 responses to LPS, included exaggerated surface temperature depression (the mice shiver and huddle together in their cages) and overt diarrhea in over 53% of the treated animals. Not surprisingly, IL-1β, the other key regulator in the fever response, is also regulated by CEACAM1 ^12^. Thus, CEACAM1 plays an inhibitory role at two levels in the fever response, and may be a candidate drug target for fever reduction.

## METHODS

### Ethics statement

This study was carried out in strict accordance with the recommendations of the Guide for the Care and Use of Laboratory Animals of the National Institutes of Health. The protocol (number 08017) was approved by the Institutional Animal Care and Use Committee (IACUC) of the City of Hope.

### Mice Strains

*Ceacam1-/-* mice were generated by Nicole Beauchemin and coworkers (McGill University, Montreal, Canada). WT C57/B6 mice were purchased from Jackson laboratory (Bar Harbor, ME). IL-6Rα (CD126)-deficient (*Il6ra*^-/-^) mice and *Stat3^flox/flox^* mice were mentioned in the publication ^72^. Mice 7–12 weeks old were used for all the experiments.

### Synthesis, radiolabeling and PET imaging of DOTH-conjugated LPS and synthesis of FAM-labeled LPS

LPS (10 mg, O55:B5 *E. coli*, Sigma-Aldrich) was made monomeric by treatment with 5 ml of 0.5% triethylamine (Sigma-Aldrich) and by sonication for 15 min on ice. After the sonication, 200 ml of LPS was removed from the solution and added to a tube containing NaIO4 (20ul,20mM, made freshly), pH 7.1. Excess NaIO4 was removed on a Zeba spin column (Thermo Scientific, IL.) after incubated 30min on ice, reacted with 26ul of DOTH (7.8mM in H2O, 202nmol), pH6.2, at RT for 2h., and then treated with 10ul of sodium cyanoborohydride (200mM in H2O, 2000nmol) at RT for 2h., followed by running a Zeba spin column again to remove excess DOTA and NaCNBH3. All reaction was protected from light ^73^. Preparation of FAM-LPS follow the protocol of the FAM conjugation from company.

### Real time RT-PCR

Total RNA was purified from cell pellets using Trizol reagent (Invitrogen) according to the manufacturer’s instructions. The concentrations and purity of extracted RNA were measured using the NanoDrop ND-1000 Spectrophotometer (NanoDrop, Wilmington, DE) demonstrating RNA with high purity (260/280 absorbance ratio between 2.1–2.2). Using Omniscript Reverse Transcription Kit (Qiagen), 1 µg of total RNA was used for the generation of cDNA as outlined by the manufacturer in a total volume of 20 µl. Following cDNA systhesis, 2 µl was used in real time RT-PCR reactions performed on CFX96 Touch Real-Time PCR Detection System (Bio-Rad) in a 20µl volume with iQ SYBR®Green Supermix (Bio-Rad), according to the manufacturer’s instructions. Primers were applied to a final concentration of 10 uM. Primer sequences are as follows: *IL-6* forward (5’-TTCCATCCAGTTGCCTTCTTGG-3’), *IL-6* reverse (5’-TTCTCATTTCCACGATTTCCCAG-3’); *TNFα* forward (5’-AGCACAGAAAGCATGATCCGC-3’), *TNFα* reverse (5’-TGCCACAAGCAGGAATGAGAAG-3’); *GAPDH* forward (5’-GTCGGTGTGAACGGATTTG-3’), *GAPDH* reverse (5’-GAACATGTAGACCATGTAGTTG-3’), *TLR4* forward (5’-ATGGCATGGCTTACACCACC-3’), *TLR4* reverse (5’-GAGGCCAATTTTGTCTCCACA-3’); *IFNβ* forward (5’-CAGCTCCAAGAAAGGACGAAC-3’), *IFNβ* reverse (5’-GGCAGTGTAACTCTTCTGCAT-3’); *IFNα* forward (5’-TGATGAGCTACTACTGGTCAGC-3’), *IFNα* reverse (5’-GATCTCTTAGCACAAGGATGGC-3’); *CCL5* forward (5’-GCTGCTTTGCCTACCTCTCC-3’), *CCL5* reverse (5’-TCGAGTGACAAACACGACTGC-3’). A TaqMan probe (Mm00462535_g1) from Life Technologies was used for detection of Regnase1. After denaturation for 3 min at 95°C, 40 cycles of amplification were performed (95°C for 10s then 55°C for 10s). Finally, melting curves were generated between 55°C and 95°C, for every 0.5°C. All Ct values were normalized to GAPDH, and quantification of gene expression was calculated by using the ΔCT method.

### Flow Cytometry and Cell Sorting

Bone marrow cells were flushed out using PBS with 2% FBS and red blood cells were lysed using red blood cell lysis buffer (Sigma-Aldrich). For cell surface staining, cells were washed with PBS, blocked with anti-mouse CD16/32 antibody, stained with antibodies described in the figures, washed 3 times with 1% BSA PBS, and analyzed with a FACSCanton II cytometer (BD Biosciences). For intracellular staining, after treated with Brefeldin A (BFA) and 500ng/mL LPS for 5 hours, cells were washed with PBS, fixed with Fixation/Permeabilization Concentrate/diluents (eBioscience, San Diego, USA), blocked with anti-mouse CD16/32 antibody, and stained with antibodies shown in the figures. After washed with PBS containing 1% BSA and 0.1% saponin, stained cells were assessed with a FACSCanton II cytometer. For cell sorting, total cells were stained with FITC conjugated lineage marker antibodies (anti-Ly-6G, anti-B220, anti-CD3, anti-Ter119, anti-NK1.1 and anti-CD19), and PE/Cy5 conjugated anti-CD135, PE/Cy7 conjugated anti-CD11b, APC conjugated anti-CD115, and APC/Cy7 conjugated anti-CD117 (Biolegend, San Diego, CA 92121) and sorted by SROP. Purity was checked using FACSCanton II and >95% purity sorted cells were used in the experiment.

### Bone marrow monocyte isolation and Cytokine cytometric bead assay

Sorted bone marrow monocytes (Mo), common monocyte progenitors (cMoP), monocyte-macrophage DC progenitors (MDP) and common DC precursors (CDP) were cultured in the concentration of 1×10^6^/mL with RPMI1640 supplement with 10% FBS and antibiotics. For surface staining, cells were stained with TLR4-PE (Clone UT41) and Isotype-PE (Clone eBR2a) (eBiosciences) and TLR4-APC (Clone SA15-21), CD14-PE (Clone Sa14-2), RP105-PE (Clone RP/14), MD1-PE (Clone MD-113) and isotypes (Biolegend, San Diego, CA 92121). For intracellular IL-6 and TNFα analysis, cells were treated with BFA and 500ng/mL LPS for 5 hours, then cells were analyzed using intracellular anti-IL-6-APC and anti-TNFα-APC (Biolegend, San Diego, CA 92121) staining. For LPS treatment or RP105 antibody treatment, cells were treated with 500ng/mL LPS or 20μg/mL anti-RP105 antibody (Clone RP/14, Biolegend) over time shown in the figures and harvested for qPCR analysis. For blocking and inhibiting experiment, cells were incubated over time shown in the figures or preincubated with TLR4 blocking antibody (Clone 76B357.1, Novus, Littleton, CO80210, USA), VAV1 inhibitor Azathioprine and 6-thio-GTP (abcam, Cambridge, MA02139, USA), CD14 blocking antibody (Clone M14-23, Biolegend), MD1 blocking antibody (Clone MD113, abeomics, San diego, CA92121), SHP-1 inhibitor PTP inhibitor III (Cayman Chemical, Ann Arbor, Michigan), Src inhibitor Src I1 (Tocris, Minneapolis, MN55413, USA), TLR4 inhibitor C34 (Tocris, Minneapolis, MN55413, USA) for 20min following treated with 500ng/mL LPS over time shown in the figures. Then, cells were harvested for qPCR or immunoblot analysis. For IL-6 release analysis, cells were treated with 500ng/mL LPS over time in the figure and supernatants were collected and analyzed using cytometric bead array. The concentration of inflammatory cytokines was measured using cytometric bead array (CBA; BD Biosciences, USA) as described previously ^74^. This assay is multiplexed and measures the concentration of each cytokine simultaneously. Mean of fluorescence intensity were converted to cytokine concentration (pg/mL) using a standard curve for each cytokine measured. Graphs were plotted using GraphPad Prism.

### Diarrheogenic activity and thermometry

The diarrheogenic activity was measured by observing wet area distance of tail from mouse anus. Any area more than 2 millimeters indicated positive diarrhea cases. Each group consisted of 17 mice and the observation lasted 48 h in all experiments. Temperatures were measured 1 hour before and 1, 2, 4, 6, 8, 24, and 48 hours after LPS i.p. injection. Temperatures were measured using non-Contact Infrared Thermometer (EXtech Instruments, Model 42505) as described ^75, 76^. The mice were manually restrained, exposing the ventral aspect of the body. Body temperature was measured by aiming the thermometer at the animal’s abdomen.

### Treatment of RAW264.7 and J774A.1 with CEACAM1 siRNA

Murine macrophage cell line RAW264.7 (ATCC® RIB-71™) and J774A.1 (ATCC® TIB-67™) were cultured for 24 hours after seeding, then transfected with CEACAM1 small interfering RNA(siRNA) or scrambled control siRNA (Origene, Rockville, MD 20850, USA) according to manufacturer’s protocol. After 24h, 48 h and 72h of transfection, cells were harvested and stained with mouse CEACAM1 APC-conjugated antibody (R&D systems, Inc., Minneapolis, MN 55413, USA) to verify the silencing effect of CEACAM1. After silencing CEACAM1 for 48 hours, CEACAM1 could not be detected on RAW264.7 and J774A.1 cells. RAW264.7 and J774A.1 cells were treated with 500ng/mL LPS over time shown in the figures or with BFA plus 500ng/mL LPS for 5 hours, then cells were harvested and analyzed using qPCR or intracellular staining. For detection of IL-6 level in the supernatant, supernatant were harvested and analyzed using cytometric bead array.

### Immunoblot analysis and immunoprecipitation (IP)

After 10mg/kg i.p. injection of LPS for 2 hours, Liver, spleen and duodenum of both WT and *Ceacam1^-/-^* mice were harvested separately, homogenized and lysed in 1% NP-40 lysis buffer as previously described ^64^. Total protein (50 µg) was separated by SDS-gel polyacrylamide electrophoresis, transferred to nitrocellulose membranes and probed with either anti-mouse phospho-gp130 (Clone A-12, Santa Cruz Biotechnology), anti-mouse phospho-STAT1, anti-mouse phospho-STAT3, anti-mouse SOCS3 or anti-β-actin antibody (Cell signaling technology, Danvers, MA 01923). Signals were detected on the Odyssey Infrared Imaging System (LI-COR Biosciences, Lincoln, NE, USA).

Bone marrow monocytes also were negatively isolated using EasySep™ Mouse Monocyte Enrichment Kit (StemCell Technologies Inc, Vancouver, Canada) according to manufacture protocol. Negatively isolated bone marrow monocytes with 1 ×10^6^ concentration in RPMI 1640 medium supplement with 10% FBS and antibiotics. After incubation in the presence or absence LPS for 15 minutes, cells were harvested and lysed on ice for 30 min, and protein concentration was determined using the Bio-Rad protein assay. Immunoprecipitation (IP) of RP105 or CEACAM1 was performed with anti-CEACAM1 (Clone MAb-CC1, Biolegend, San Diego, CA) or RP105 (Clone RP/14) using Pierce protein A/G plus agarose (Thermo Scientific, Rockford, IL) per the manufacturer’s protocol, immunoblotted with appropriate primary antibodies (anti-MD1 pAb from Santa Cruz Biotechnology, Inc., Dallas, TX.; anti-RP105 pAb and anti-CD14 pAb from Abcam, Cambridge, MA; phospho-VAV1 Y160 pAb from Bioss Antibodies Inc., Woburn, MA; VAV1 pAb, SHP1 pAb, phospho-SHP-1 Y564 pAb, Src, and 4G10 pAb from Cell Signaling Technology, Inc., Danvers, MA; anti-β-actin mAb from GeneTex, Inc., Irvine, CA) and infrared-labeled IRDye secondary antibodies. Detection was carried out using the Odyssey infrared imaging ^77^.

### Statistical Analysis

Assay results were expressed as means ±SEM and paired or unpaired Student’s t-tests were used for comparisons. All p-values are two-sided. Data were analyzed with GraphPad Prism software (version 5.0, GraphPad Software, San Diego, CA, USA).

## ACKNOWLEDGEMENTS

The authors thank Dr. Angel Gu, Frances Chang, Courtni Salinas and Jennifer Chean in our laboratory and Dr. Walter Tsark in animal facility for technical support. The authors thank flow cytometry core for cell sorting and small animal imaging core for PET imaging.

## CORRESPONDING AUTHORS

Correspondence to Zhifang Zhang or John E. Shively.

## CAPTIONS FOR SUPPORTING INFORMATION

**Fig S1. *IL-6* mRNA expression in peritoneal cavity tissues after LPS challenge.**

**Fig S2. Intracellular IL-6 and TNFα staining of hepatocytes and Kupffer cells in response to LPS in *Ceacam1^-/-^* mice.**

**Fig S3. IL-6 and TNFα levels of liver cells after LPS treatment in vitro and IL-6 receptor downstream signaling activation after i.p. LPS in vivo**

**Fig S4. CEACAM1 expression on bone marrow CD115^+^ (M-CSF^+^) cells and CD115^+^ cell pattern change of bone marrow cells after treated with LPS + BFA.**

**Fig S5. CEACAM1 expression on bone marrow CD115^+^ (M-CSF^+^) cells and CD115^+^ cell pattern change of bone marrow cells after treated with LPS + BFA.**

**Fig S6. RAW264.7 cells start to produce IL-6 after treatment with LPS+BFA for 11 hour, while silencing of CEACAM1 does not affect IL-6 production in murine macrophage RAW264.7 cells at the 2 hour and 24 hour time points.**

**Fig S7. Murine macrophage cell line J774A.1 does not produce IL-6 within 5 hours after LPS treatment, while silencing of CEACAM1 does not interfere with IL-6 production after LPS treatment at 2 hour and 24 hour points.**

**Fig S8. PTP inhibitor III, a SHP1 inhibitor, increases levels of phospho-VAV1 in bone marrow monocytes of WT mice.**

**Fig S9. CEACAM1, TLR4 and RP105 expression on human peripheral blood monocytes.**

